# Metagenome assembled genomes of novel taxa from an acid mine drainage environment

**DOI:** 10.1101/2020.07.02.185728

**Authors:** Christen L. Grettenberger, Trinity L. Hamilton

**Affiliations:** Department of Earth and Planetary Sciences, University of California Davis; Department of Plant and Microbial Biology and The Biotechnology Institute, University of Minnesota, St. Paul, MN 55108

**Author notes:** For correspondence: Christen L. Grettenberger 2119 Earth and Physical Sciences Building, University of California Davis, Davis USA, 95776, Trinity L Hamilton. 218 Cargill Building, Plant and Microbial Biology, University of Minnesota, St. Paul, USA, 55108.

**Keywords:** AMD, metagenome assembled genome, biogeochemical cycling, bioremediation

## Abstract

Acid mine drainage (AMD) is a global problem in which iron sulfide minerals oxidize and generate acidic, metal-rich water. Bioremediation relies on understanding how microbial communities inhabiting an AMD site contribute to biogeochemical cycling. A number of studies have reported community composition in AMD sites from16S rRNA gene amplicons but it remains difficult to link taxa to function, especially in the absence of closely related cultured species or those with published genomes. Unfortunately, there is a paucity of genomes and cultured taxa from AMD environments. Here, we report 29 novel metagenome assembled genomes from Cabin Branch, an AMD site in the Daniel Boone National Forest, KY, USA. The genomes span 11 bacterial phyla and include one Archaea and include taxa that contribute to carbon, nitrogen, sulfur, and iron cycling. These data reveal overlooked taxa that contribute to carbon fixation in AMD sites as well as uncharacterized Fe(II)-oxidizing bacteria. These data provide additional context for 16S rRNA gene studies, add to our understanding of the taxa involved in biogeochemical cycling in AMD environments, and can inform bioremediation strategies.

**IMPORTANCE:** Bioremediating acid mine drainage requires understanding how microbial communities influence geochemical cycling of iron and sulfur and biologically important elements like carbon and nitrogen. Research in this area has provided an abundance of 16S rRNA gene amplicon data. However, linking these data to metabolisms is difficult because many AMD taxa are uncultured or lack published genomes. Here, we present metagenome assembled genomes from 29 novel AMD taxa and detail their metabolic potential. These data provide information on AMD taxa that could be important for bioremediation strategies including taxa that are involved in cycling iron, sulfur, carbon, and nitrogen.

## MAIN TEXT

### 1. Introduction

Acid mine drainage (AMD) is a global environmental problem. Oxidative processes, both biotic and abiotic, release protons and reduced metals from sulfide minerals, resulting in highly acidic and toxic conditions that degrade environmental quality. Due to the toxicity and environmental impact of AMD, bioremediation strategies have become of interest. Research in AMD environments often seeks to understand the biogeochemical cycling occurring in the environment and aims to inform and improve the bioremediation of these sites (1–4). The geochemistry at these sites relies on the microbial communities inhabiting them. Biotic oxidation of reduced metal sulfides contributes to the formation of AMD while sulfate and iron reduction can both decrease the concentration of soluble metals and increase pH (5–11). However, the metabolic potential of many taxa in AMD environments remains uncharacterized because these taxa are not closely related to cultured taxa or those with published genomes.

AMD environments are characterized by redox gradients including contrasting concentration of oxygen and reduced metals. They can also vary in heavy metal content and pH. However, 16S rRNA gene surveys reveal that many of the same species inhabit AMD sites across the globe. For example, *Ferrovum* spp. are found in the Appalachian Coal Belt (12–14), the Iberian Pyrite Belt (15, 16), Wales (17, 18), and southeast and southwest China (19, 20). Therefore, the importance of these groups is not limited to a single site, and lack of information about their metabolisms hinders investigations of AMD environments world-wide. For example, Archaea within the order Thermoplasmatales are commonly found in AMD sites worldwide, especially those with low pH (9, 21–23). However, in many instances, 16S rRNA sequences isolated from AMD are only distantly related to cultured Thermoplasmatales. Taxa in the Thermoplasmatales perform diverse metabolisms, including Fe(II) oxidation (24), obligate heterotrophy (25, 26), and sulfur respiration (27). Therefore, it is difficult to infer their metabolic potential (22). Similarly, taxa within newly discovered phyla like the Elusimicrobiota (formerly Termite Group 1) and Eremiobacteriota (formerly the WPS-2) inhabit AMD sites (28, 29), but these groups have few if any cultured taxa. Given the widespread distribution of these lineages, these taxa may play an important role in biogeochemical cycling in AMD environments, but without closely-related cultured relatives or well-annotated genomes, it is not possible to elucidate their role or potential use in bioremediation strategies.

Even in well-studied AMD groups like the Gammaproteobacteria, multiple closely related taxa may occur in AMD sites but may play different roles from their close relatives. For example, multiple *Ferrovum* taxa differ in their ability to fix nitrogen (30–33). This intragenus metabolic diversity complicates our ability to understand biogeochemical cycling in AMD environments.

Obtaining a species in pure culture has long been considered the gold standard for determining the biogeochemical role that a taxon may play in the environment. However, characterized isolates from AMD environments are rare. Culturing taxa is inherently time-consuming, especially those that require micro-oxic conditions, and can be difficult because species often require co-occurring taxa. For example, in culture, *Ferrovum* sp. co-occur with heterotrophic organisms that remove pyruvic acid and other organic material (34). Metagenomic sequencing has proven to be a valuable tool for guiding isolation of common AMD microbes through the recovery of near complete genomes. Tyson et al. used a genome-directed approach to isolate a *Leptospirillum ferrooxidans* spp. capable of nitrogen fixation (35). Metagenomic approaches also provide valuable information about community structure and diversity. Thus, ‘omics-based approaches can complement pure culture studies, provide valuable insight to biogeochemical cycling in AMD environments, and inform bioremediation strategies in the absence of fully characterized isolates.

Here, we present 29 novel, high-quality, metagenome-assembled genomes (MAGs) from Cabin Branch, an acid mine drainage site in the Daniel Boone National Forest, KY, USA. These data suggest AMD environments host previously uncharacterized Fe(II)-oxidizing bacteria and highlight the metabolic potential of a number of microbes commonly recovered in 16S rRNA-based studies of AMD. These genomes will provide additional context for gene amplicon studies in AMD environments, aid in culturing these taxa in the future, and could inform AMD bioremediation strategies.

### 2. Methods

#### 2.1 Site location

Cabin Branch is an acid mine drainage site in the Daniel Boone National Forest in Kentucky, near the border with Tennessee. Groundwater flows out from an emergence and across the limestone-lined channel before entering a pond, the Rose Pool. The microbial communities within Cabin Branch are dominated by the Fe(II)-oxidizing taxon *Ferrovum myxofaciens* (13). Methods for sample collection, DNA extraction and sequencing, and metagenome assembly and binning were described previously (33) and we include brief descriptions below.

#### 2.2 Molecular Analyses

##### 2.2.1 Sample Collection, DNA Extraction, and Sequencing

Triplicate samples from each site were collected for DNA extraction and were flash frozen and stored at −80 °C until processed. DNA was extracted from each replicate sample using a DNeasy PowerSoil Kit (Qiagen, Carlsbad, CA, USA) and quantified using a Qubit 3.0 Fluorometer (Invitrogen, Burlington, ON, Canada). Extractions were pooled and submitted to the University of Minnesota Genomics Center for metagenomic sequencing and sequenced using HiSeq2500 High-Output 2 x 125 bp chemistry. Three samples were sequenced per lane.

##### 2.2.2. Metagenomic analysis

Trimmed, quality-controlled sequences were assembled using MegaHit (36) using standard parameters except minimum contig length, which was set at 1000 base pairs. Reads were mapped to the assembly using bowtie2 (37) and depth was calculated using the jgi_summarize_bam_contig_depths command in Anvi’o v. 6.1 (38). Binning was performed in MetaBAT using default parameters (39) and CheckM was used to determine bin completeness (40). Bins >70% complete with <3% contamination were selected for further analysis. The average nucleotide identity (ANI) across the surviving bins was calculated using anvi-compute-ani in Anvi’o v. 6.1 which uses the PyANI algorithm to compute ANI (38, 41). Bins that shared >99% ANI across the genome were considered to be the same taxon. For each taxon, the bin with the highest completion was selected for further analysis. Bins were uploaded to KBASE and annotated using the “annotate assembly and re-annotate genomes with prokka” app (v. 1.12(42) and classified using the GTDB app (43–46).

Single copy, ribosomal protein sequences from Campbell et al., were retrieved from the MAGs and reference genomes, concatenated, and aligned in Anvi’o (47). Anvi’o uses muscle to align the concatenated sequences (48). Maximum likelihood trees were constructed using RAxML-HPC2 on XSEDE in the CIPRES Science Gateway using standard parameters: 100 bootstrap iterations, a Protein CAT model, DAYHOFF protein substitution matrix, and no correction for ascertainment bias (49, 50). Trees were visualized and rooted in the interactive tree of life (51). Newick formatted tree files are available in the supplementary information.

Relative abundance of each MAG was determined by mapping reads from each metagenome against each MAG using BBMap (52). The pileup tool within BBMap was used to summarize mapped read and the relative abundance was calculated from the total number of mapped reads divided by the total number of reads in the metagenome (53). Unmapped reads and reads mapping to more than one region were removed using SAMtools (54) prior to pileup.

Metabolic pathways for carbon, nitrogen, and sulfur cycling were predicted in each MAG using METABOLIC v. 3.0 (55).

Cyc2 genes were identified using BLAST to identify genes homologous to the Cyc2 like cytochrome c involved in Fe(II) oxidation (56–58). A BLAST database was constructed using the Cyc2 retrieved previously (57) and the search was performed using an e-value of 1E-5. To ensure that the sequences retrived with the BLAST search were homologous to the Cyc2-like protein involved in Fe(II) oxidation, retrieved sequences and those described previously (57) were aligned in MAFFT and a maximum likelihood tree was constructed as described above. Quality-controlled, unassembled, metagenomic data are available in the NCBI Sequence Read Archive under accession numbers SRR9677580 - SRR9677585. The metagenome assembled genomes used for analysis are also available on NCBI under accession numbers SAMN14771053 to SAMN14771081.

### Results

Cabin Branch is an AMD site in the Daniel Boone National Forest in southern Kentucky, US. Groundwater at Cabin Branch emerges at pH 2.90 and flows down a limestone-lined channel with a pH of 2.92 (installed as a passive remediation strategy) and enters a pool (“Rose Pool) which has a pH of 2.97. Dissolved oxygen increases down the drainage site (77.5 µmol/L at the emergence, 401 µmol/L in Rose Pool) and Fe(II) concentration ranges from 403 µmol/L at the emergence, 882.0 µmol/L in the limestone-lined channel, and 11.16 µmol/L in Rose Pool (33)

We recovered 256 bins from metagenomes: 38 from the emergence, 32 from the limestone-lined channel, 66 from the Rose Pool, and 120 from the coassembly. Of these, 56 were >70% complete with <3% contamination. These bins belonged to 32 unique taxa (Table 1). Here we present 29 novel, high-quality, metagenome-assembled genomes (MAGs): 4 from the emergence, 7 from the limestone-lined channel, 9 from Rose Pool and 9 from the co-assembled data. The MAGs ranged in relative abundance from ∼4.3% to ∼0.17% (Figures 1 and 2). The *Ferrovum* MAGs (MAG 23 and and MAG24) were described in (33)and MAG 7 is closely related to a previously described genome (59, 60).

**Table 1.**
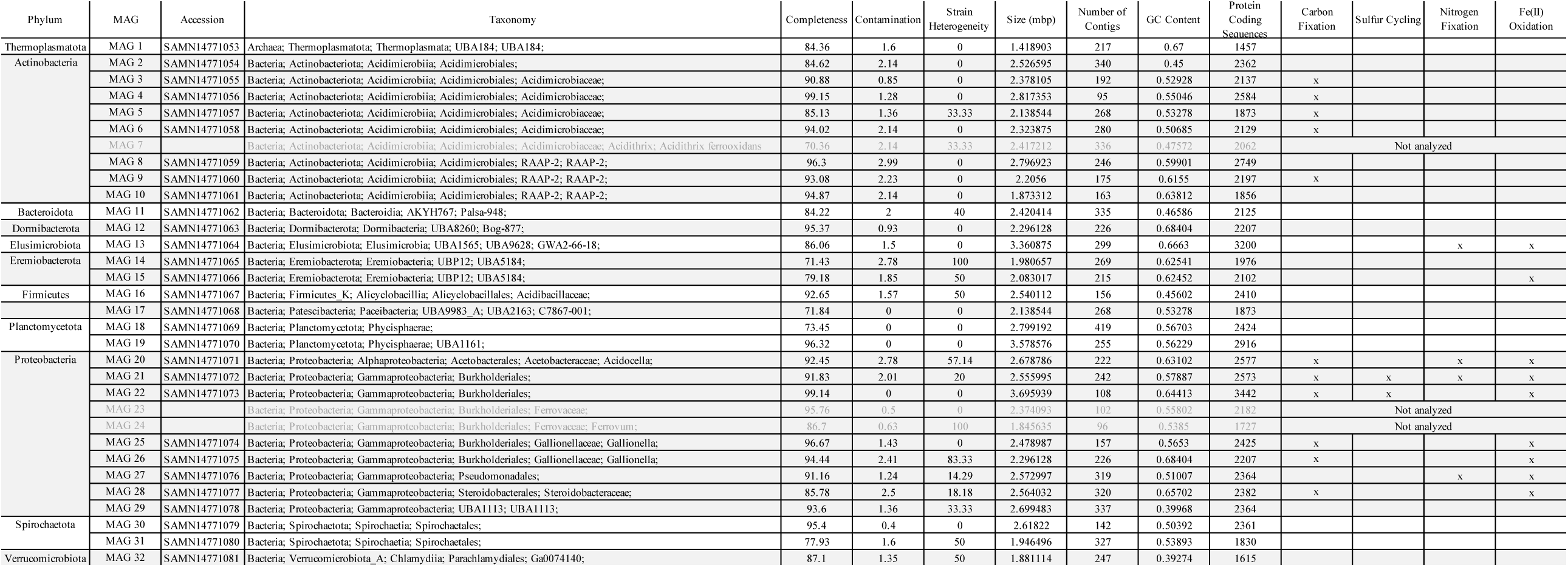
Summary of the MAGs presented here. Taxa indicated by grey text were not analyzed in this work because they were either closely related to cultured taxa (e.g. MAG 7) or were presented previously (MAGs 24 and 24; 33).

**Figure 1.**
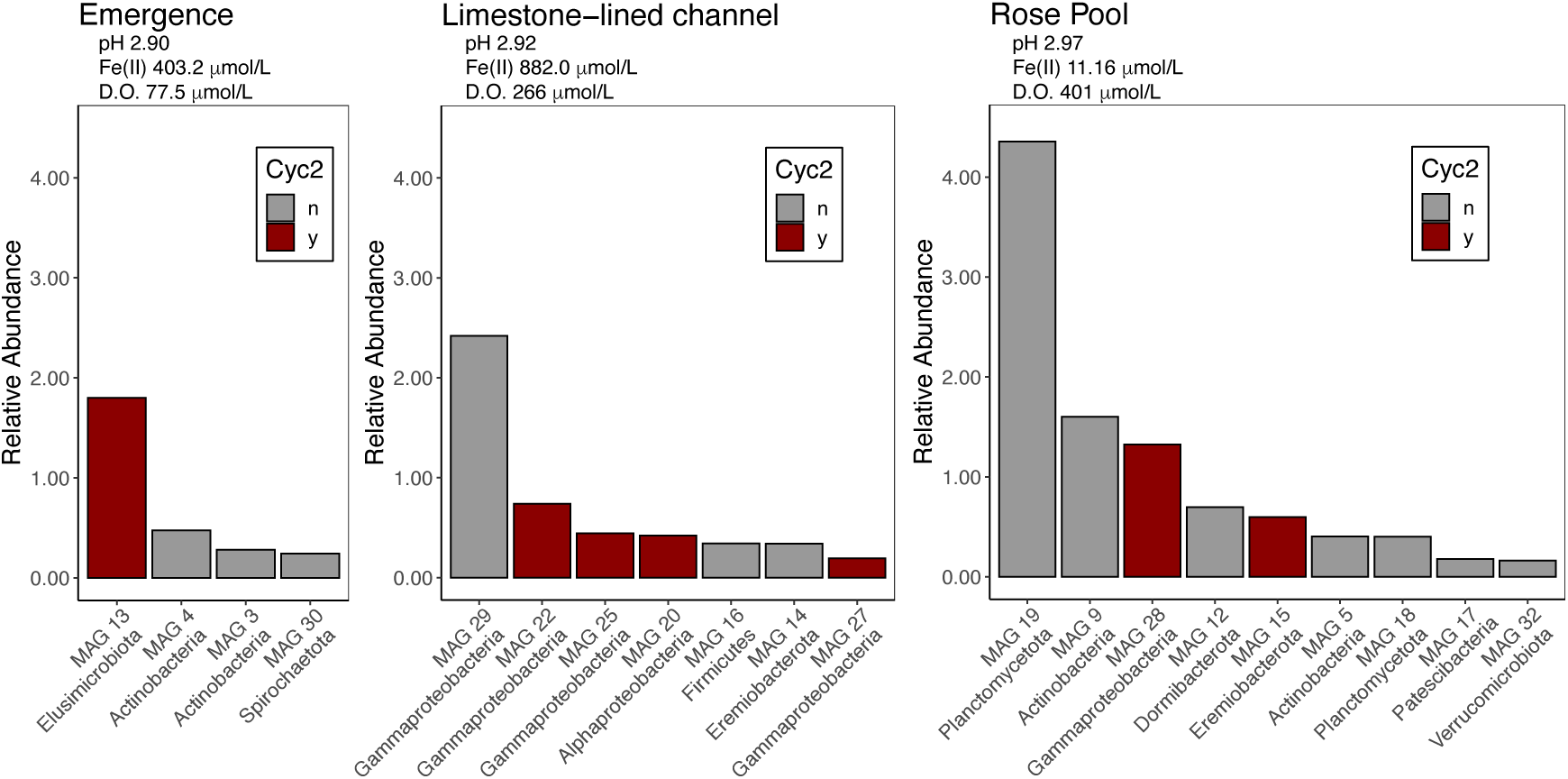
Rank abundance curve of the relative abundance of MAGs recovered from the emergence, the limestone-lined channel and Rose Pool. pH, Fe(II), and D.O. were measured at the time of sample collection and are reported in (33). Red bars indicate MAGs that encode Cyc2. D.O., dissolved oxygen.

**Figure 2.**
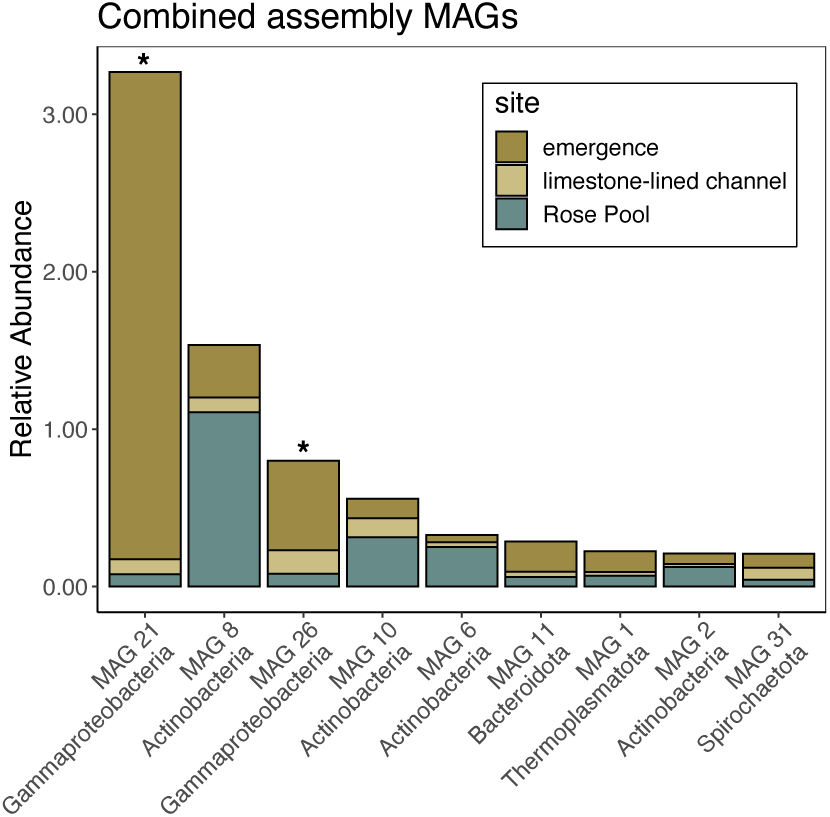
Rank abundance curve of the relative abundance of MAGs recovered from the co-assembled data in each site — the emergence, the limestone-lined channel and Rose Pool. Asterisks denote MAGs that encode Cyc2.

Below, we examine functions that are most relevant to AMD ecosystems including aerobic respiration, carbon fixation, nitrogen cycling and biogeochemical cycling of sulfur and iron in each MAG by phylogenetic group. For sulfur cycling, we focus on dissimilatory sulfate reduction and sulfur oxidation by examining the presence or absence of *dsr* and *sox* genes. The metabolic potential for ferrous iron oxidation was based on the presence of *Cyc2* like genes that may be involved in this process (56, 57). None of the MAGs contain complete genomes and a gene that is absent in the MAG may be present in the taxon. Therefore, these data indicate the genes present in, not absent from, a taxon. A summary of these taxa is available in Table 1. More complete genome descriptions and the output from METABOLIC are available in the supplemental material.

#### 3.1. Archaea

We recovered a single MAG classified as Archaea (MAG 1). The MAG belonged to the *Thermoplasmata* and was most closely related to *Methanomassiliicoccus* spp. (Figure 1). It did not encode genes associated with carbon fixation, N_2_ fixation, Fe(II) oxidation or dissimilatory sulfur cycling. It appears to be capable of heterotrophic metabolisms including the degradation of some aromatics and acetogenesis. MAG 1 was recovered from the co-assembly and was most abundant in the emergence (Figure 2).

#### 3.2. Bacteria

##### 3.2.1. Actinobacteria

We retrieved nine actinobacterial MAGs, all of which belong to the order Acidimicrobiales (Figure 3). MAGs 3 – 7 belonged to the Acidimicrobiaceae, but MAGs 3-6 were unidentified below this level. MAGs 8 - 10 were affiliated with the family and genus RAAP-2 but were unidentified at the species level. Five taxa (MAGs 3 – 6, and 9) encode genes for carbon fixation. Actinobacteria MAGs were recovered from the emergence and Rose Pool (Figure 1) as well as the co-assembly (Figure 2).

**Figure 3.**
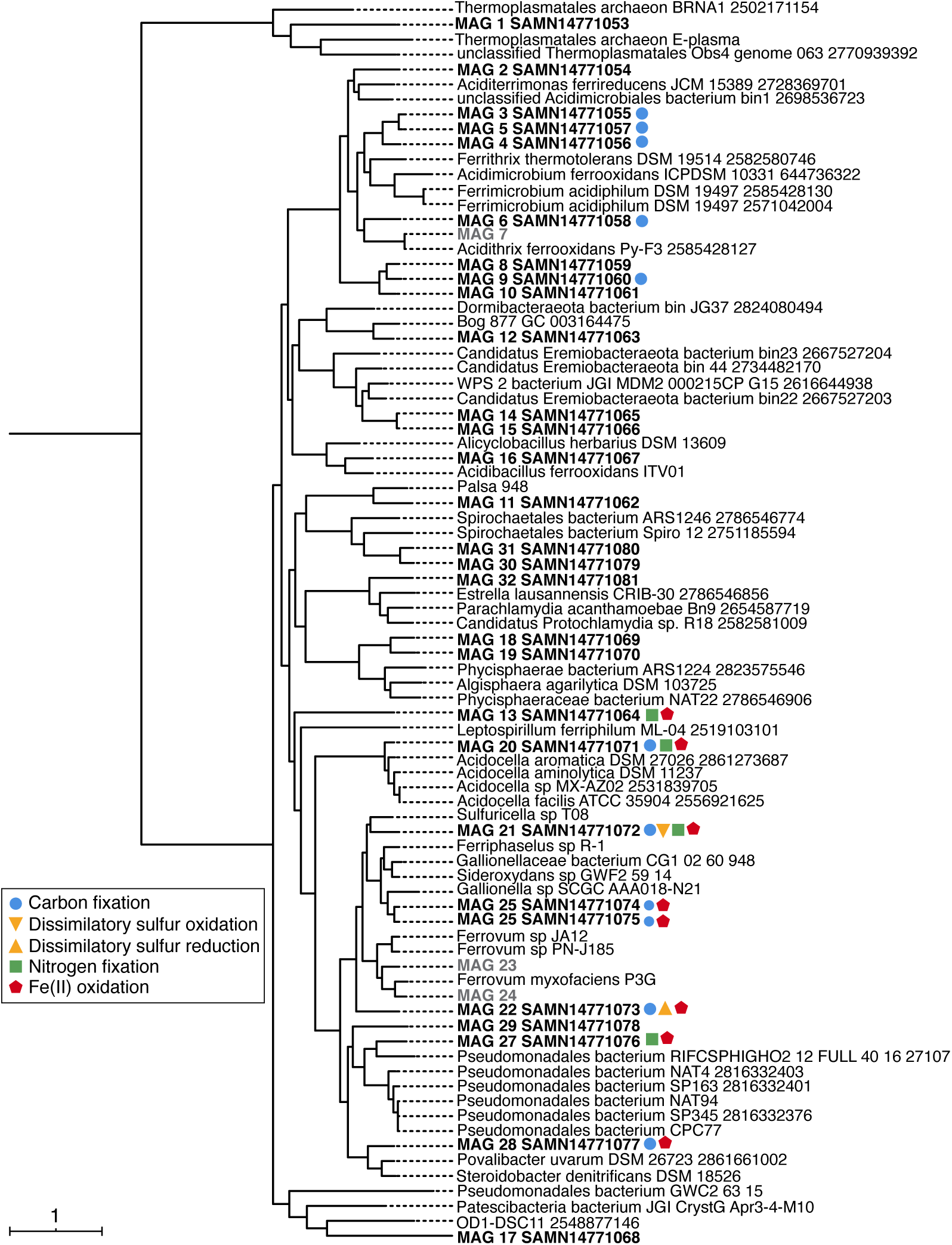
Concatenated single-copy marker gene tree constructed using genes from (47). The tree contains 83 taxa. MAGs from Cabin Branch are indicated in bold. Shapes indicated the metabolic potential of the MAGs. Carbon fixation in blue circles. Dissimilatory sulfur oxidation in yellow triangles with points up and reduction in triangles with points down, nitrogen fixation in green squares, and Fe(II) oxidation in red stars.

##### 3.2.2. Bacteroidota

We retrieved a single taxon from the phylum Bacterioidota (MAG 11) from the co-assembly (Figures 2 and 3). It was affiliated with the class Bacteroidia, order AKH767, and family Palsa-948, and was unclassified below this level. The most closely related taxon was retrieved from thawing permafrost (61). It encodes the genes necessary for nitrous oxide reduction.

##### 3.2.3. Dormibacteraeota

We retrieved one taxon from the phylum Dormibacteraeota (MAG 12). This taxon was affiliated with the class Dormibacteria, the order UBA8260, and the family Bog-877 and is most closely related to a taxon from thawing permafrost (61). It did not encode genes for carbon fixation, N_2_ fixation, denitrification, dissimilatory sulfur cycling, or Fe(II) oxidation. It appears to be capable of some fermentative and C1 metabolisms (File S1)

##### 3.2.4. Elusimicrobiota

We retrieved a single taxon, MAG 13, from the phylum Elusimicrobiota (formerly Termite Group 1). It was affiliated with the class Elusimicrobia, order UBA1565, family UBA9628, and genus GWA2-66-18. This taxon is most closely related to one from an aquifer system (62). It encodes the genes necessary for N_2_ fixation and nitric oxide reduction. It also appears to encode for a Cyc2 like cytochrome that it may use for Fe(II) oxidation. MAG 13 was recovered from the emergence with a relative abundance of ∼1.8%.

##### 3.2.5. Eremiobacteriota

We retrieved two taxa, MAGs 14 and 15, within the phylum Eremiobacteriota (formerly the WPS-2). Both taxa were affiliated with the class Eremiobacteria, order UPB12, and family UBA5184. Neither bin contain genes for carbon fixation, N_2_ fixation, denitrification, dissimilatory sulfur cycling, or Fe(II) oxidation. Both taxa appear to be capable of heterotrophic metabolisms including aromatics degradation and acetogenesis. MAG 14 was recovered from the limestone-lined channel while MAG 15 was present in Rose Pool (Figure 1).

##### 3.2.6. Firmicutes

We retrieved a single taxon from the phylum Firmicutes (MAG 16) from the limestone-lined channel (Figure 1). This taxon was affiliated with the class Alicyclobacillia, the order Alicyclobacillales, and the family Acidibacillaceae. This taxon was most closely related to *Acidobacillus ferrooxidans*, a Fe(II) and sulfide mineral oxidizing species isolated from an AMD environments (Figure 1) (63). Unlike its closest relative, it does not encode the genes necessary for Fe(II) or sulfide mineral oxidation.

##### 3.2.7. Patescibacteria

We retrieved a single taxon, MAG 17, from the phylum Patescibacteria (formerly Candidate Phylum Radiation) from Rose Pool (Figure 1). This taxon was affiliated with the class Paceibacteria, the order UBA9983_A, the family UBA2163, and the genus C7867-001. It does not encode any of the genes of interest.

##### 3.2.8. Planctomycetota

We recovered two taxa from the phylum Planctomycetota (MAGs 18 and 19). Both taxa were affiliated with the class Phycisphaerae. One, MAG 18, was unclassified below this level. The other, MAG 19, was affiliated with the order UBA1161. Both MAG 18 and 19 were recovered from Rose Pool where MAG 19 was more abundant (4.3% and 0.4% respectively, Figure 1).

##### 3.2.9. Proteobacteria

We recovered ten taxa from the Proteobacteria. One, MAG 20, is a member of the Alphaproteobacteria. Nine are from the gammaproteobacterial order Burkholderiales. Two of these were described previously and were not analyzed here (33). Three encode genes for N_2_ fixation (MAGs 20, 21, and 27), six encode genes for carbon fixation, (MAGs 20 – 22, 25, 26, 28), seven encode genes for Fe(II) oxidation (MAGs 20-22 and 25-28), and three encode genes for partial sulfate reduction (MAGs 22, 25, and 26). Gammaproteobacterial MAGs were recovered from the limestone-lined channel, Rose Pool, and the co-assembly (Figure 1 and 2). MAG 21 from the co-assembly was particularly abundant at the emergence (3.3%) while MAG 29 was abundant in the limestone-lined channel (2.4%).

##### 3.2.10. Spirochaetota

We recovered two taxa from the phylum Spirochaetota (MAGs 30 and 31) from Rose Pool (Figure 1). MAG 30 was recovered from the emergence and MAG 31 was present in the co-assembly with low relative abundance across all sites (Figures 1 and 2). Both bins were members of the class Spirochaetia and the order Spirochaetales. One encodes the genes necessary for nitrate reduction to ammonia.

##### 3.2.11. Verrucomicrobiota

We recovered a single taxon from the Verrucomicrobiota (MAG 32). It contains some of the genes necessary for aerobic respiration and acetogenesis.

### 4. Discussion

#### 4.1 Carbon Fixation

Lithotrophic carbon fixation can be a significant source of primary productivity in AMD ecosystems (64). At Cabin Branch, *Ferrovum* spp. are abundant, ranging from 5 - 33%, and likely contribute to primary productivity (33). Here, we recovered eleven MAGs that encode the genes necessary for carbon fixation. These autotrophs include those that are closely related to known lithoautotrophic organisms, including *Gallionella* (MAGs 25 and 26) and other Bulkholderiales (e.g. MAGs 20-22) as well as heterotrophs including *Acidocella* (e.g. MAG 20). This indicates that primary productivity in AMD sites may be driven, in part, by organisms that have not been considered in the past.

#### 4.2 Sulfur Cycling

Sulfur cycling is an important process in AMD ecosystems. Bioremediation may rely on dissimilatory sulfate reduction, especially in constructed wetlands (4). Dissimilatory sulfate reduction combats AMD by generating alkalinity, can lead to the formation of ferrous sulfide minerals in sediments, and decreases the concentration of soluble metals (5–11). Conversely, biological sulfur oxidation generates AMD by oxidizing sulfur in iron sulfide minerals. MAG 21 encodes homologs of *dsrA, dsrB*, and *aprA*, indicating that it may be capable of dissimilatory sulfate reduction, at least from APS to sulfide. Therefore, this taxon may play an important role in constructed wetland bioremediation.

AMD occurs naturally when weathering processes expose sulfide mineral-bearing rocks to oxygen-rich water. The result is the oxidation of these sulfide minerals which produces sulfuric acid (H_2_SO_4_) and dissolved metals. Iron-sulfide minerals like pyrite can also be oxidized by biological sulfur oxidation (65, 66). The oxidation of iron sulfide minerals may occur either at a pyrite vein or in freshly deposited sediments that are exposed to oxygen. We recovered a single MAG, MAG 22, that contains the genes necessary for sulfur oxidation from thiosulfate to sulfate. This taxon is unlikely to cause the oxidation of sulfide-bearing minerals but may play a role in aqueous sulfur cycling in the environment.

#### 4.3 Nitrogen Cycling

Common Fe(II) oxidizing organisms in AMD environments such as *Ferrovum myxofaciens* are capable of nitrogen fixation (30–33, 67) and may provide fixed nitrogen to AMD communities. Four of the MAGs recovered here (MAGS 13, 20, 21, and 27) contain the genes necessary for nitrogen fixation. These organisms may serve as a source of bioavailable nitrogen in AMD ecosystems and, in so doing, increase the productivity of their communities.

#### 4.4. Fe(II) Oxidation

Fe(II) oxidation is a key process in AMD environments for bioremediation and as a source of energy to drive primary productivity. Indeed, our previous analyses recovered multiple abundant Ferrovum MAGs whose genomes are consistent with carbon fixation couples to Fe oxidation (33). Here we identified 9 MAGs that encoded homologs of the Cyc2 protein involved in Fe(II) oxidation (57). These MAGs were recovered across sample sites that range in dissolved oxygen and Fe(II) concentration and are present at relative abundances that suggest key roles in community function (Figures 1 and 2). Seven of the MAGs that encode Cyc2 belong to proteobacterial lineages with other known Fe oxidizers. Additionally, MAGs within the recently discovered phyla Elusimicrobia (MAG 13) and Eremiobacterota (MAG 15) encode Cyc2. These taxa have not previously been recognized as Fe(II) oxidizers but the recovery of Cyc2 in these MAGs further expands our knowledge of the taxonomic diversity of Fe(II) oxidation.

#### 4.5 Phylogenetic Relatedness and Metabolism

In the absence of characterized isolates, we often rely on phylogenetic relationships between taxa found at AMD sites and their closest cultured relatives to infer their role in biogeochemical cycling (13, 22, 68, 69). This approach leverages the use of 16S rRNA gene amplicon data, which is relatively inexpensive in terms of time and cost, at the expense of the metabolic insights inferred from expensive and time-consuming ‘omics approaches or validated by culture-based approaches. This approach—inferring physiology from 16 rRNA gene sequences—can be informative for major metabolic pathways when taxa are closely related to their nearest cultured relative. For example, like *Gallionella ferruginea*, the metabolic potential of the *Gallionella* MAGs (MAGs 25 and 26) recovered here is consistent with chemolithoautotrophy fueled by Fe(II) oxidation. These relationships are less robust with increasing phylogenetic distance.

Inferring metabolism from16S rRNA gene sequences becomes more difficult as the number of available genomes from similar environments decreases. For example, AMD environments often host organisms within the Archaeal order Thermaplasmatales (9, 22, 23). However, there is a paucity of Thermaplasmatales genomes available from AMD environments. This order also contains taxa with diverse metabolisms, including Fe(II) oxidation (24), obligate heterotrophy (25, 26), and sulfur respiration (27). The lack of genomes and culture representatives from AMD environments coupled to the physiological diversity of Thermaplasmatales makes it difficult to interpret the role of these archaea AMD environments. The Thermoplasmatales MAG presented here appears to be a heterotroph capable of aerobic respiration.

The role of taxa affiliated with uncultivated or recently discovered phyla in biogeochemical cycling in AMD is particularly difficult to predict. Here, we presented MAGs from four such phyla — the Dormibacterota, the Elusimicrobiota, the Eremiobacterota, and the Patescibacteria. Dormibacterota and Patescibacteria are not widely reported in AMD. Eremiobacterota inhabit multiple mining-impacted sites including stalactites in a mining cave (70), neutral mine drainage in Brazil (71), and AMD in the eastern United States (28). Abundances of the Eremiobacterota correlated with total organic carbon in an AMD site in China (68) and were recovered from Rose Pool were dissolved organic carbon is present (36.8 d from the emergence where dissolved oxygen is present, albeit well below saturation (77.5 µmol/L, (33). One of the Eremiobacterota MAGs (MAG 15) One of the MAGs at Cabin Branch encodes a Cyc-2 like protein that may be involved in Fe(II) oxidation. Therefore, this taxon may play an important and underappreciated role in Fe cycling in AMD environments.

Elusimicrobiota have also been found in AMD environments across the globe including in Spain (70, 72), France (29), and Svalbard (73), but it is not abundant in these environments. The only cultivated taxa from this phylum are strictly anaerobic (74–76). The Elusimicrobiota MAG from Cabin Branch encodes genes for three terminal oxidases and likely employs aerobic respiration. The Elusimicrobiota MAG (MAG 13) was abundant in the emergence where dissolved oxygen is present, albeit below saturation (77.5 µmol/L). The MAG also contains the genes necessary for nitrogen fixation and may encode a Cyc2-like protein that it may use for Fe(II) oxidation. Therefore, it likely plays an important role in nitrogen cycling in AMD environments and may also contribute to iron cycling.

A combination of ‘omics-based approaches and cultivation can increase our ability to correlate function with taxonomy from 16S rRNA amplicon studies. Here we present high quality MAGs from AMD sites to increase current understanding of community composition and function. These data reveal previously unrecognized taxa that contribute to carbon, nitrogen, and Fe(II) cycling in AMD. In particular, these data underscore roles for previously uncharacterized Gammaproteobacteria in Fe(II) oxidation in addition to uncultivated or recently discovered phyla, the prevalence of Actinobacteria across AMD sites that range in oxygen and Fe(II) concentration, and taxa with high relative abundance whose function remains unclear. These data provide a framework to assist in culturing taxa of interest as well as additional target organisms for AMD bioremediation strategies.

## ACKNOWLEDGEMENTS

We are grateful to the staff of the National Forest Service and Daniel Boone National Forest, especially Margueritte Wilson and Claudia Cotton, for the advice and insight regarding mine locations. We thank A. Gangidine, M. Berberich, R. Jain, and C. Schuler for assistance in field sampling and processing samples in the laboratory. The authors acknowledge the Minnesota Supercomputing Institute (MSI) at the University of Minnesota for providing resources that contributed to the research results reported within this paper.

## AUTHOR CONTRIBUTIONS

C.L.G and T.L.H designed the study, completed the analyses and wrote the paper.

## MATERIALS & CORRESPONDENCE

Correspondence and requests for materials should be addressed to T.L.H.: Trinity L. Hamilton. Department of Plant and Microbial Biology, University of Minnesota, St. Paul, USA, 55108. Phone: +16126256372.

## SUPPLEMENTAL RESULTS

Descriptions of each MAG retrieved from the Cabin Branch AMD site.

## SUPPLEMENTAL FILES

**File S1**. Output from METABOLIC showing presence or absence of the genes necessary for metabolic pathways for each MAG.

**File S2**. Newick-formatted, maximum likelihood tree of concatenated ribosomal proteins for all MAGs.

**File S3**. Newick-formatted, maximum likelihood tree of Cyc2-like proteins retrieved from Cabin Branch MAGs and reference sequences.

## REFERENCE CITED

1. Gupta A, Dutta A, Sarkar J, Panigrahi MK, Sar P. 2018. Low-Abundance Members of the Firmicutes Facilitate Bioremediation of Soil Impacted by Highly Acidic Mine Drainage From the Malanjkhand Copper Project, India. Front Microbiol 9:2882.

2. Johnson DB, Hallberg KB. 2005. Biogeochemistry of the compost bioreactor components of a composite acid mine drainage passive remediation system. Sci Total Environ 338:81–93.

3. Neculita C-M, Zagury GJ, Bussière B. 2007. Passive Treatment of Acid Mine Drainage in Bioreactors using Sulfate-Reducing Bacteria. J Environ Qual 36:1–16.

4. Sánchez-Andrea I, Sanz JL, Bijmans MFM, Stams AJM. 2013. Sulfate reduction at low pH to remediate acid mine drainage. J Hazard Mater 269:98–109.

5. Baker BJ, Banfield JF. 2003. Microbial communities in acid mine drainage. Fems Microbiol Ecol 44:139–152.

6. Bijmans MFM, Vries E de, Yang C-H, Buisman CJN, Lens PNL, Dopson M. 2010. Sulfate reduction at pH 4.0 for treatment of process and wastewaters. Biotechnol Progr 26:1029–37.

7. Bijmans MFM, Helvoort P-J van, Dar SA, Dopson M, Lens PNL, Buisman CJN. 2009. Selective recovery of nickel over iron from a nickel–iron solution using microbial sulfate reduction in a gas-lift bioreactor. Water Res 43:853–861.

8. Church CD, Wilkin RT, Alpers CN, Rye RO, McCleskey RB. 2007. Microbial sulfate reduction and metal attenuation in pH 4 acid mine water. Geochem T 8:10.

9. Druschel GK, Baker BJ, Gihring TM, Banfield JF. 2004. Acid mine drainage biogeochemistry at Iron Mountain, California. Geochem T 5:13.

10. Giloteaux L, Duran R, Casiot C, Bruneel O, Elbaz-Poulichet F, Goñi-Urriza M. 2012. Three-year survey of sulfate-reducing bacteria community structure in Carnoulès acid mine drainage (France), highly contaminated by arsenic. Fems Microbiol Ecol 83:724–737.

11. Kaksonen AH, Franzmann PD, Puhakka JA. 2004. Effects of hydraulic retention time and sulfide toxicity on ethanol and acetate oxidation in sulfate-reducing metal-precipitating fluidized-bed reactor. Biotechnol Bioeng 86:332–343.

12. Grettenberger CL, Pearce AR, Bibby KJ, Jones DS, Burgos WD, Macalady JL. 2017. Efficient Low-pH Iron Removal by a Microbial Iron Oxide Mound Ecosystem at Scalp Level Run. Appl Environ Microb 83:e00015–17.

13. Havig JR, Grettenberger C, Hamilton TL. 2017. Geochemistry and microbial community composition across a range of acid mine drainage impact and implications for the Neoarchean-Paleoproterozoic transition. J Geophys Res Biogeosciences 122:1404–1422.

14. Jones DS, Kohl C, Grettenberger C, Larson LN, Burgos WD, Macalady JL. 2015. Geochemical Niches of Iron-Oxidizing Acidophiles in Acidic Coal Mine Drainage. Appl Environ Microb 81:1242–1250.

15. González-Toril E, Santofimia E, López-Pamo E, Omoregie EO, Amils R, Aguilera Á. 2013. Microbial Ecology in Extreme Acidic Pit Lakes from the Iberian Pyrite Belt (SW Spain). Adv Mat Res 825:23–27.

16. Santofimia E, González-Toril E, López-Pamo E, Gomariz M, Amils R, Aguilera A. 2013. Microbial Diversity and Its Relationship to Physicochemical Characteristics of the Water in Two Extreme Acidic Pit Lakes from the Iberian Pyrite Belt (SW Spain). Plos One 8:e66746.

17. Hallberg KB, Coupland K, Kimura S, Johnson DB. 2006. Macroscopic Streamer Growths in Acidic, Metal-Rich Mine Waters in North Wales Consist of Novel and Remarkably Simple Bacterial Communities. Appl Environ Microb 72:2022–2030.

18. Kay C, Rowe O, Rocchetti L, Coupland K, Hallberg K, Johnson D. 2013. Evolution of Microbial “Streamer” Growths in an Acidic, Metal-Contaminated Stream Draining an Abandoned Underground Copper Mine. Life 3:189–210.

19. Kuang J-L, Huang L-N, Chen L-X, Hua Z-S, Li S-J, Hu M, Li J-T, Shu W-S. 2013. Contemporary environmental variation determines microbial diversity patterns in acid mine drainage. Isme J 7:1038.

20. Sun W, Xiao T, Sun M, Dong Y, Ning Z, Xiao E, Tang S, Li J. 2015. Diversity of the Sediment Microbial Community in the Aha Watershed (Southwest China) in Response to Acid Mine Drainage Pollution Gradients. Appl Environ Microb 81:4874–84.

21. González-Toril E, Aguilera A, Souza-Egipsy V, Ercilla MD, López-Pamo E, Sánchez-España FJ, Amils R. 2009. Comparison between Acid Mine Effluents, La Zarza-Perrunal and Río Tinto (Iberian Pyritic Belt). Adv Mat Res 71–73:113–116.

22. Grettenberger CL, Rench RLM, Gruen DS, Mills DB, Carney C, Brainard J, Hamasaki H, Ramirez R, Watanabe Y, Amaral-Zettler LA, Ohmoto H, Macalady JL. 2020. Microbial population structure in a stratified, acidic pit lake in the Iberian Pyrite Belt. Geomicrobiol J 1–12.

23. Qiu G, Wan M, Qian L, Huang Z, Liu K, Liu X, Shi W, Yang Y. 2008. Archaeal diversity in acid mine drainage from Dabaoshan Mine, China. J Basic Microb 48:401–409.

24. Golyshina OV, Pivovarova TA, Karavaiko GI, Kondratéva TF, Moore ER, Abraham WR, Lünsdorf H, Timmis KN, Yakimov MM, Golyshin PN. 2000. Ferroplasma acidiphilum gen. nov., sp. nov., an acidophilic, autotrophic, ferrous-iron-oxidizing, cell-wall-lacking, mesophilic member of the Ferroplasmaceae fam. nov., comprising a distinct lineage of the Archaea. Int J Syst Evol Micr 50:997–1006.

25. Golyshina OV, Lünsdorf H, Kublanov IV, Goldenstein NI, Hinrichs K-U, Golyshin PN. 2015. The novel extremely acidophilic, cell-wall-deficient archaeon Cuniculiplasma divulgatum gen. nov., sp. nov. represents a new family, Cuniculiplasmataceae fam. nov., of the order Thermoplasmatales. Int J Syst Evol Micr 66:332–40.

26. Itoh T, Yoshikawa N, Takashina T. 2007. Thermogymnomonas acidicola gen. nov., sp. nov., a novel thermoacidophilic, cell wall-less archaeon in the order Thermoplasmatales, isolated from a solfataric soil in Hakone, Japan. Int J Syst Evol Micr 57:2557–2561.

27. Segerer A, Langworthy TA, Stetter KO. 1988. Thermoplasma acidophilum and Thermoplasma volcanium sp. nov. from Solfatara Fields. Syst Appl Microbiol 10:161–171.

28. Brantner JS, Haake ZJ, Burwick JE, Menge CM, Hotchkiss ST, Senko JM. 2014. Depthdependent geochemical and microbiological gradients in Fe(III) deposits resulting from coal mine-derived acid mine drainage. Front Microbiol 5:215.

29. Volant A, Bruneel O, Desoeuvre A, Héry M, Casiot C, Bru N, Delpoux S, Fahy A, Javerliat F, Bouchez O, Duran R, Bertin PN, Elbaz-Poulichet F, Lauga B. 2014. Diversity and spatiotemporal dynamics of bacterial communities: physicochemical and other drivers along an acid mine drainage. Fems Microbiol Ecol 90:247–263.

30. Ullrich SR, Poehlein A, Daniel R, Tischler JS, Vogel S, Schlömann M, Mühling M. 2015. Comparative Genomics Underlines the Functional and Taxonomic Diversity of Novel “*Ferrovum*” Related Iron Oxidizing Bacteria. Adv Mat Res 1130:15–18.

31. Ullrich SR, González C, Poehlein A, Tischler JS, Daniel R, Schlömann M, Holmes DS, Mühling M. 2016. Gene Loss and Horizontal Gene Transfer Contributed to the Genome Evolution of the Extreme Acidophile “Ferrovum.” Front Microbiol 7:797.

32. Ullrich SR, Poehlein A, Tischler JS, González C, Ossandon FJ, Daniel R, Holmes DS, Schlömann M, Mühling M. 2016. Genome Analysis of the Biotechnologically Relevant Acidophilic Iron Oxidising Strain JA12 Indicates Phylogenetic and Metabolic Diversity within the Novel Genus “Ferrovum.” Plos One 11:e0146832.

33. Grettenberger CL, Havig J, Hamilton T. n.d. Metabolic diversity and co-occurrence of multiple Ferrovum species at an acid mine drainage site.

34. Johnson DB, Hallberg KB, Hedrich S. 2014. Uncovering a Microbial Enigma: Isolation and Characterization of the Streamer-Generating, Iron-Oxidizing, Acidophilic Bacterium “Ferrovum myxofaciens.” Appl Environ Microb 80:672–680.

35. Tyson GW, Lo I, Baker BJ, Allen EE, Hugenholtz P, Banfield JF. 2005. Genome-Directed Isolation of the Key Nitrogen Fixer Leptospirillum ferrodiazotrophum sp. nov. from an Acidophilic Microbial Community. Appl Environ Microb 71:6319–6324.

36. Li D, Liu C-M, Luo R, Sadakane K, Lam T-W. 2015. MEGAHIT: an ultra-fast single-node solution for large and complex metagenomics assembly via succinct de Bruijn graph. Bioinformatics 31:1674–1676.

37. Langmead B, Salzberg SL. 2012. Fast gapped-read alignment with Bowtie 2. Nat Methods 9:357.

38. Eren AM, Esen ÖC, Quince C, Vineis JH, Morrison HG, Sogin ML, Delmont TO. 2015. Anvi’o: an advanced analysis and visualization platform for ‘omics data. Peerj 3:e1319.

39. Kang DD, Froula J, Egan R, Wang Z. 2015. MetaBAT, an efficient tool for accurately reconstructing single genomes from complex microbial communities. Peerj 3:e1165.

40. Parks DH, Imelfort M, Skennerton CT, Hugenholtz P, Tyson GW. 2015. CheckM: assessing the quality of microbial genomes recovered from isolates, single cells, and metagenomes. Genome Res 25:1043–1055.

41. Pritchard L, Glover RH, Humphris S, Elphinstone JG, Toth IK. 2016. Genomics and taxonomy in diagnostics for food security: soft-rotting enterobacterial plant pathogens. Anal Methods-uk 8:12–24.

42. Seemann T. 2014. Prokka: rapid prokaryotic genome annotation. Bioinformatics 30:2068–2069.

43. Arkin AP, Cottingham RW, Henry CS, Harris NL, Stevens RL, Maslov S, Dehal P, Ware D, Perez F, Canon S, Sneddon MW, Henderson ML, Riehl WJ, Murphy-Olson D, Chan SY, Kamimura RT, Kumari S, Drake MM, Brettin TS, Glass EM, Chivian D, Gunter D, Weston DJ, Allen BH, Baumohl J, Best AA, Bowen B, Brenner SE, Bun CC, Chandonia J-M, Chia J-M, Colasanti R, Conrad N, Davis JJ, Davison BH, DeJongh M, Devoid S, Dietrich E, Dubchak I, Edirisinghe JN, Fang G, Faria JP, Frybarger PM, Gerlach W, Gerstein M, Greiner A, Gurtowski J, Haun HL, He F, Jain R, Joachimiak MP, Keegan KP, Kondo S, Kumar V, Land ML, Meyer F, Mills M, Novichkov PS, Oh T, Olsen GJ, Olson R, Parrello B, Pasternak S, Pearson E, Poon SS, Price GA, Ramakrishnan S, Ranjan P, Ronald PC, Schatz MC, Seaver SMD, Shukla M, Sutormin RA, Syed MH, Thomason J, Tintle NL, Wang D, Xia F, Yoo H, Yoo S, Yu D. 2018. KBase: The United States Department of Energy Systems Biology Knowledgebase. Nat Biotechnol 36:566.

44. Allen B, Drake M, Harris N, Sullivan T. 2017. Using KBase to Assemble and Annotate Prokaryotic Genomes. Curr Protoc Microbiol 46:1E.13.1-1E.13.18.

45. Parks DH, Chuvochina M, Waite DW, Rinke C, Skarshewski A, Chaumeil P-A, Hugenholtz P. 2018. A standardized bacterial taxonomy based on genome phylogeny substantially revises the tree of life. Nat Biotechnol 36:996–1004.

46. Chaumeil P-A, Mussig AJ, Hugenholtz P, Parks DH. 2019. GTDB-Tk: a toolkit to classify genomes with the Genome Taxonomy Database. Bioinform Oxf Engl.

47. Campbell BJ, Yu L, Heidelberg JF, Kirchman DL. 2011. Activity of abundant and rare bacteria in a coastal ocean. Proc National Acad Sci 108:12776–12781.

48. Edgar RC. 2004. MUSCLE: multiple sequence alignment with high accuracy and high throughput. Nucleic Acids Res 32:1792–1797.

49. Miller MA, Pfeiffer W, Schwartz T. 2010. Creating the CIPRES Science Gateway for Inference of Large Phylogenetic Trees. 2010 Gatew Comput Environ Work Gce 1–8.

50. Stamatakis A. 2006. RAxML-VI-HPC: maximum likelihood-based phylogenetic analyses with thousands of taxa and mixed models. Bioinformatics 22:2688–2690.

51. Letunic I, Bork P. 2016. Interactive tree of life (iTOL) v3: an online tool for the display and annotation of phylogenetic and other trees. Nucleic Acids Res 44:W242–W245.

52. Bushnell B. 2014. BBMap: A fast, accurate, splice-aware aligner.

53. Hua Z-S, Wang Y-L, Evans PN, Qu Y-N, Goh KM, Rao Y-Z, Qi Y-L, Li Y-X, Huang M-J, Jiao J-Y, Chen Y-T, Mao Y-P, Shu W-S, Hozzein W, Hedlund BP, Tyson GW, Zhang T, Li W-J. 2019. Insights into the ecological roles and evolution of methyl-coenzyme M reductase-containing hot spring Archaea. Nat Commun 10:4574.

54. Li H, Handsaker B, Wysoker A, Fennell T, Ruan J, Homer N, Marth G, Abecasis G, Durbin R, Subgroup 1000 Genome Project Data Processing. 2009. The Sequence Alignment/Map format and SAMtools. Bioinform Oxf Engl 25:2078–9.

55. Zhou Z, Tran P, Liu Y, Kieft K, Anantharaman K. 2019. METABOLIC: A scalable high-throughput metabolic and biogeochemical functional trait profiler based on microbial genomes. Biorxiv 761643.

56. Chan C, McAllister SM, Garber A, Hallahan BJ, Rozovsky S. 2018. Fe oxidation by a fused cytochrome-porin common to diverse Fe-oxidizing bacteria. Biorxiv 228056.

57. McAllister SM, Polson SW, Butterfield DA, Glazer BT, Sylvan JB, Chan CS. 2020. Validating the Cyc2 Neutrophilic Iron Oxidation Pathway Using Meta-omics of Zetaproteobacteria Iron Mats at Marine Hydrothermal Vents. Msystems 5.

58. Altschul SF, Gish W, Miller W, Myers EW, Lipman DJ. 1990. Basic local alignment search tool. J Mol Biol 215:403–410.

59. Jones RM, Johnson DB. 2015. Acidithrix ferrooxidans gen. nov., sp. nov.; a filamentous and obligately heterotrophic, acidophilic member of the Actinobacteria that catalyzes dissimilatory oxido-reduction of iron. Res Microbiol 166:111–120.

60. Eisen S, Poehlein A, Johnson DB, Daniel R, Schlömann M, Mühling M. 2015. Genome Sequence of the Acidophilic Ferrous Iron-Oxidizing Isolate Acidithrix ferrooxidans Strain Py-F3, the Proposed Type Strain of the Novel Actinobacterial Genus Acidithrix. Genome Announc 3:e00382–15.

61. Woodcroft BJ, Singleton CM, Boyd JA, Evans PN, Emerson JB, Zayed AAF, Hoelzle RD, Lamberton TO, McCalley CK, Hodgkins SB, Wilson RM, Purvine SO, Nicora CD, Li C, Frolking S, Chanton JP, Crill PM, Saleska SR, Rich VI, Tyson GW. 2018. Genome-centric view of carbon processing in thawing permafrost. Nature 560:49–54.

62. Anantharaman K, Brown CT, Hug LA, Sharon I, Castelle CJ, Probst AJ, Thomas BC, Singh A, Wilkins MJ, Karaoz U, Brodie EL, Williams KH, Hubbard SS, Banfield JF. 2016. Thousands of microbial genomes shed light on interconnected biogeochemical processes in an aquifer system. Nat Commun 7:13219.

63. Dall’Agnol H, Ñancucheo I, Johnson DB, Oliveira R, Leite L, Pylro VS, Holanda R, Grail B, Carvalho N, Nunes GL, Tzotzos G, Fernandes GR, Dutra J, Orellana SC, Oliveira G. 2016. Draft Genome Sequence of “Acidibacillus ferrooxidans” ITV01, a Novel Acidophilic Firmicute Isolated from a Chalcopyrite Mine Drainage Site in Brazil. Genome Announc 4:e01748–15.

64. Havig JR, Hamilton TL. 2019. Productivity and Community Composition of Low Biomass/High Silica Precipitation Hot Springs: A Possible Window to Earth’s Early Biosphere? Life Basel Switz 9:64.

65. Nordstrom DK, Southam G. 1997. Geomicrobiology of sulfide mineral oxidation, p. 361–390. In Banfield, JF and Nealson, KH eds., Geomicrobioogy: Interactions between microbes and minerals, vol 35, Reviews in Mineralogy, Min Soc Am, Washington DC.

66. Evangelou VP (Bill), Zhang YL. 1995. A review: Pyrite oxidation mechanisms and acid mine drainage prevention. Crit Rev Env Sci Tec 25:141–199.

67. Moya-Beltrán A, Cárdenas JP, Covarrubias PC, Issotta F, Ossandon FJ, Grail BM, Holmes DS, Quatrini R, Johnson DB. 2014. Draft Genome Sequence of the Nominated Type Strain of “Ferrovum myxofaciens,” an Acidophilic, Iron-Oxidizing Betaproteobacterium. Genome Announc 2:e00834–14.

68. Sun W, Xiao E, Krumins V, Dong Y, Xiao T, Ning Z, Chen H, Xiao Q. 2016. Characterization of the microbial community composition and the distribution of Fe-metabolizing bacteria in a creek contaminated by acid mine drainage. Appl Microbiol Biot 100:8523–8535.

69. Senko JM, Wanjugi P, Lucas M, Bruns MA, Burgos WD. 2008. Characterization of Fe(II) oxidizing bacterial activities and communities at two acidic Appalachian coalmine drainage-impacted sites. Isme J 2:1134.

70. Méndez-García C, Mesa V, Sprenger RR, Richter M, Diez MS, Solano J, Bargiela R, Golyshina OV, Manteca Á, Ramos JL, Gallego JR, Llorente I, Santos VAPM dos, Jensen ON, Peláez AI, Sánchez J, Ferrer M. 2014. Microbial stratification in low pH oxic and suboxic macroscopic growths along an acid mine drainage. Isme J 8:1259–74.

71. Pereira LB, Vicentini R, Ottoboni LMM. 2014. Changes in the Bacterial Community of Soil from a Neutral Mine Drainage Channel. Plos One 9:e96605.

72. Mesa V, Gallego JLR, González-Gil R, Lauga B, Sánchez J, Méndez-García C, Peláez AI. 2017. Bacterial, Archaeal, and Eukaryotic Diversity across Distinct Microhabitats in an Acid Mine Drainage. Front Microbiol 8:1756.

73. García-Moyano A, Austnes AE, Lanzén A, González-Toril E, Aguilera Á, Øvreås L. 2015. Novel and Unexpected Microbial Diversity in Acid Mine Drainage in Svalbard (78° N), Revealed by Culture-Independent Approaches. Microorg 3:667–94.

74. Geissinger O, Herlemann DPR, Mörschel E, Maier UG, Brune A. 2009. The ultramicrobacterium “Elusimicrobium minutum” gen. nov., sp. nov., the first cultivated representative of the termite group 1 phylum. Appl Environ Microb 75:2831–40.

75. Herlemann DPR, Geissinger O, Ikeda-Ohtsubo W, Kunin V, Sun H, Lapidus A, Hugenholtz P, Brune A. 2009. Genomic analysis of “Elusimicrobium minutum,” the first cultivated representative of the phylum “Elusimicrobia” (formerly termite group 1). Appl Environ Microb 75:2841–9.

76. Zheng H, Dietrich C, Radek R, Brune A. 2015. E ndomicrobium proavitum, the first isolate of E ndomicrobia class. nov. (phylum E lusimicrobia) - an ultramicrobacterium with an unusual cell cycle that fixes nitrogen with a Group IV nitrogenase: Endomicrobium proavitum gen. nov. sp. nov. Environ Microbiol 18:191–204.

